## Supplementary figures and images for "SPAG7 deletion causes intrauterine growth restriction, resulting in adulthood obesity and metabolic dysfunction"

### Supplemental Figures

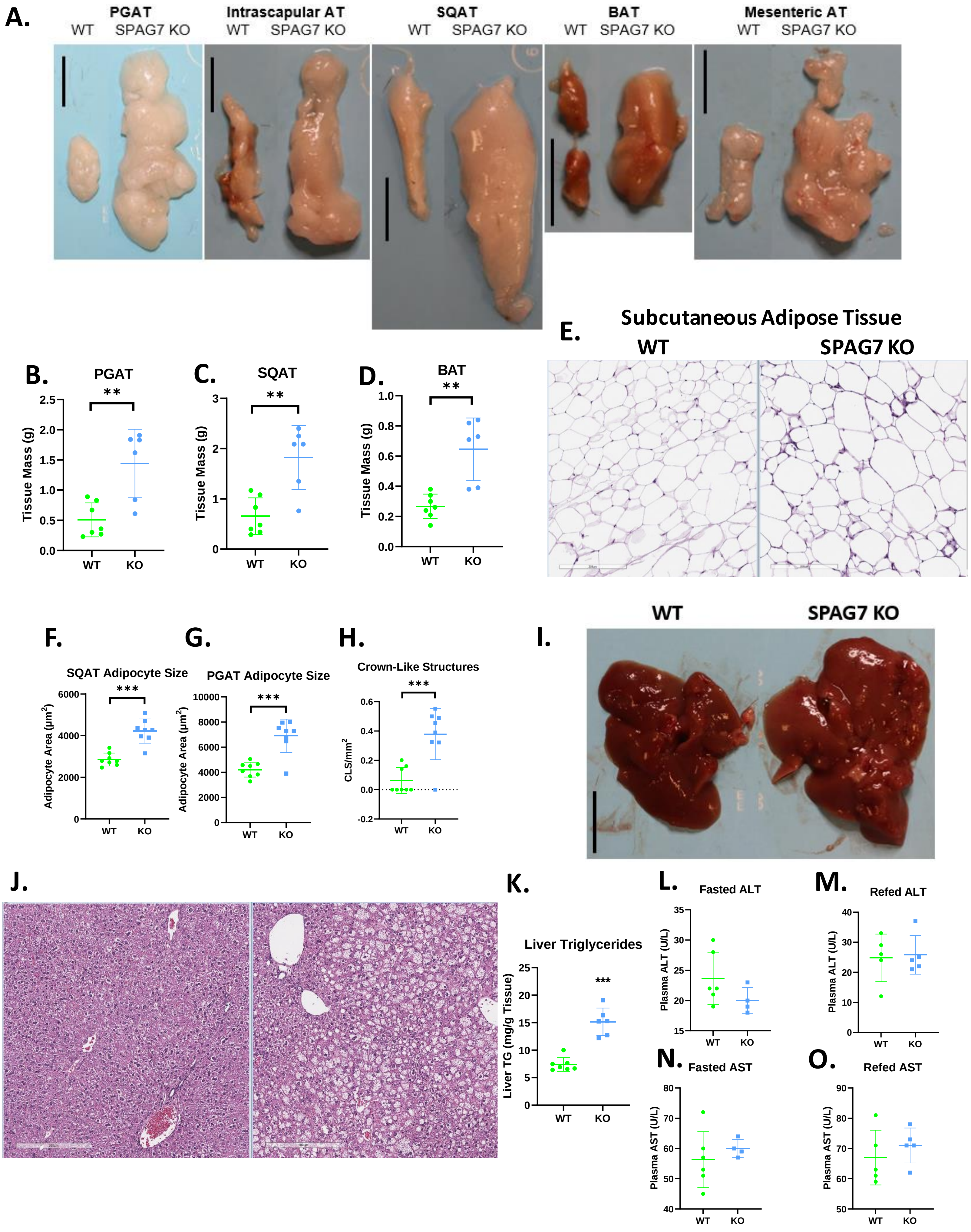

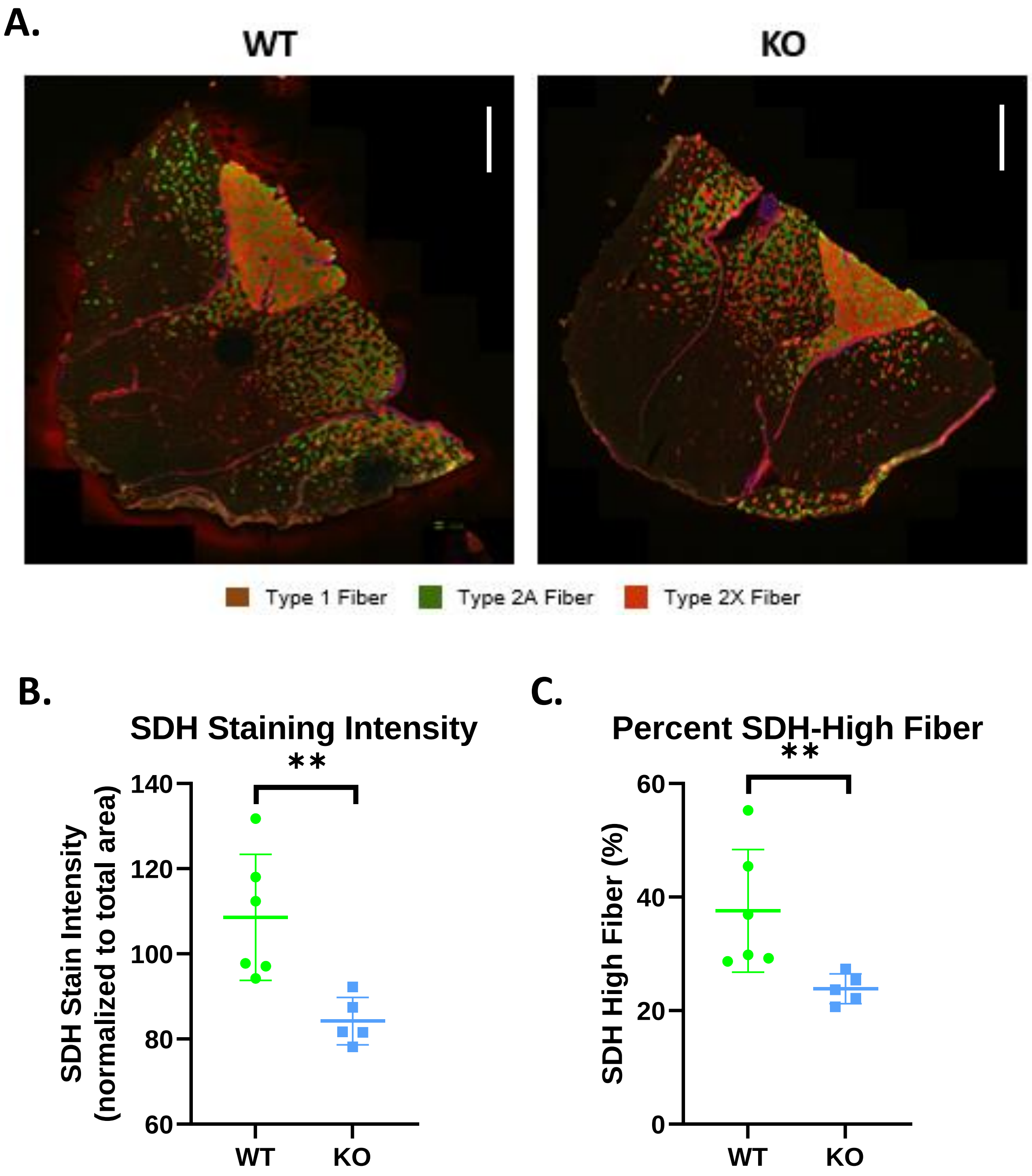

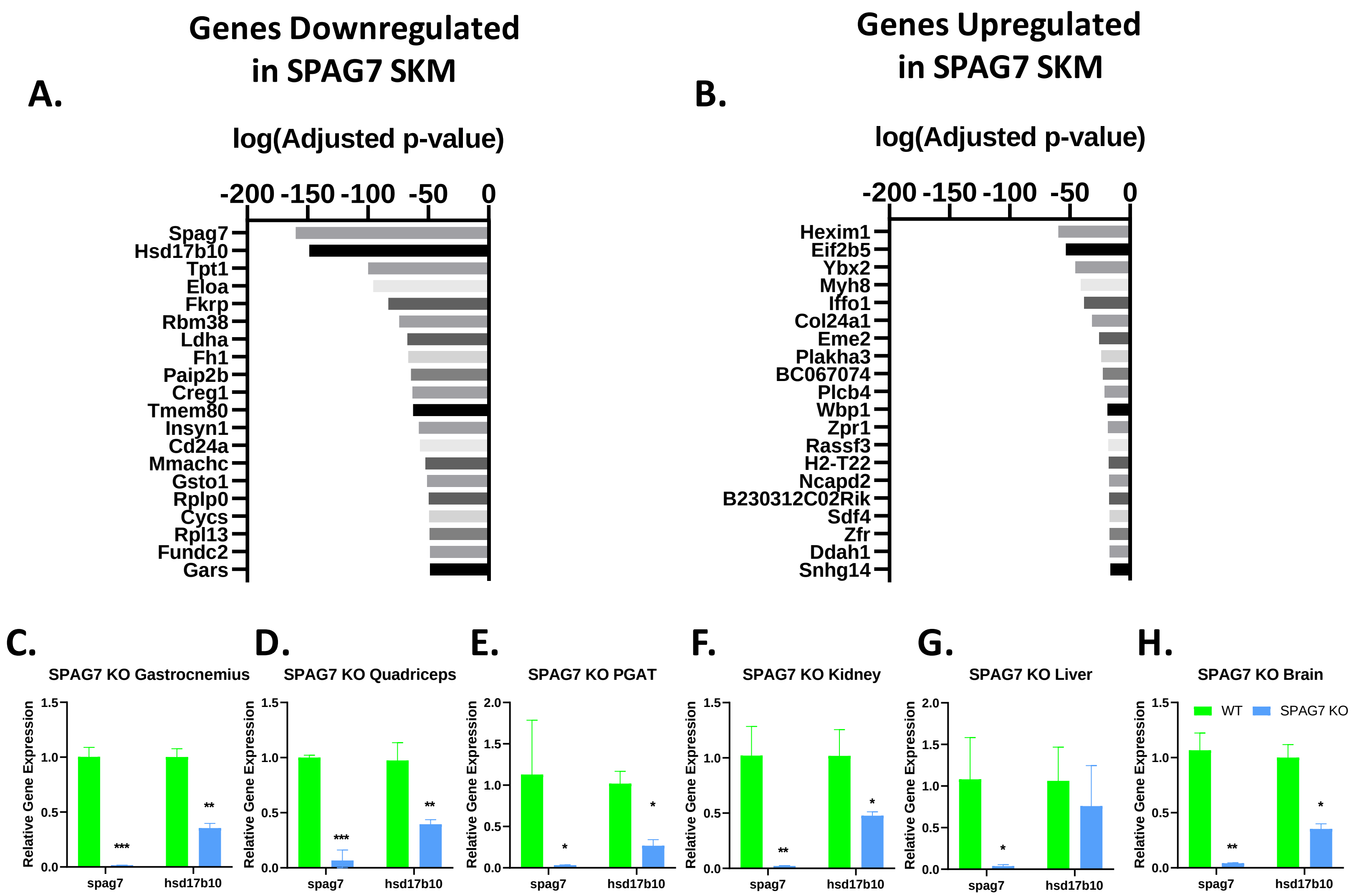
